# Spatial working memory enhances gamma oscillations in rodent olfactory bulb and medial prefrontal cortex

**DOI:** 10.64898/2026.10.13.758724

**Authors:** Morteza Mooziri, Ali Samii Moghaddam, Mohammad Reza Raoufy, Mohammad Ali Mirshekar

## Abstract

Spatial working memory (SWM), a specialized form of working memory that temporarily encodes and retains spatial information, is fundamental to guide animal behaviors, yet our current knowledge of its circuit mechanisms remains far from known. Since, olfactory bulb (OB) and medial prefrontal cortex (mPFC) are critical brain structures for a wide range of cognitive behaviors, we hypothesized that the OB-mPFC circuit gamma oscillations might support rodent SWM processing. Our results show that SWM enhances regional low (30-60 Hz), but not high (60-100 Hz), gamma oscillations in both OB and mPFC, in correlation with the animals behavioral performance. Additionally, the OB-mPFC gamma band functional connectivity is strengthened by SWM. Moreover, we find that mPFC gamma oscillatory information transfer to OB supports the animals SWM performance. We also show that OB low gamma oscillations are modulated by mPFC delta phase, which further signifies the roles of mPFC feedbacks to shape OB gamma activity during SWM. These results suggest that OB-mPFC circuit gamma rhythms underlie SWM processing, and point towards possible mechanisms for cognitive disorders.

## 1. Introduction

Working memory (WM), the ability to temporarily maintain and manipulate information, is fundamental to mammalian higher-order cognition [1-5], and its disruption are observed across neuropsychiatric disorders [6, 7]. Spatial WM (SWM), a specialized form of WM that encodes and retains spatial locations over short delays, is essential for fundamental behaviors such as navigation [8]. Despite decades of research, the neural mechanisms underlying SWM—particularly the contribution of distributed brain circuits and their dynamic interactions—remain incompletely understood. Elucidating the oscillatory and connectivity signatures of SWM is therefore critical for understanding both normal cognition and its pathological disruptions.

The medial prefrontal cortex (mPFC) has long been recognized as a central hub for WM, integrating sensory inputs, maintaining representations during delay periods, and guiding behavioral responses [1-3, 5, 9-17]. In parallel, the olfactory bulb (OB), traditionally viewed as a primary sensory relay [18-20], is increasingly implicated in cognitive and emotional processes beyond olfaction [12, 21-29]. Furthermore, allergic diseases of the respiratory system impose behavioral disorders along with neural activity disruptions in OB, mPFC, and other brain structures [14, 22, 30-32]. Importantly, interactions between OB and mPFC oscillations support a wide range of behaviors, including SWM [21, 25, 26, 28, 29], raising the possibility that the OB–mPFC circuit plays a broader role in cognitive control and SWM.

Neural oscillations, particularly in the gamma frequency range (> 30 Hz), are thought to underlie/coordinate information processing within/across brain regions [16, 25, 26, 33-35]. Gamma oscillations are further subdivided into slower and faster sub-bands, which may reflect distinct computational processes [10, 30, 36, 37]. In the context of working memory, gamma-band activity in prefrontal cortex has been associated with the active maintenance and reactivation of representations, as well as the coordination of distributed neuronal ensembles [16, 33, 35, 38, 39]. Similarly, gamma oscillations in sensory regions are modulated by behavioral state and task demands [33, 40, 41]. Furthermore, task-relevant enhancement of gamma power during WM has been reported in multiple brain structures [1, 38, 39], suggesting that gamma rhythms might serve as a common mechanism for this purpose. Importantly, in the OB–mPFC circuit, despite established roles for the slow oscillations [17], there is lack of evidence on how faster neural activities are modulated by SWM.

Beyond local oscillatory dynamics, inter-regional communication is also shown to be essential for SWM, with particular examples in the OB–mPFC circuit [12, 14, 25, 26]. Abundance of anatomical and functional connections between OB and mPFC facilitates such long-range interactions [21, 22, 25, 28, 29, 42]. For instance, rodent SWM has been associated with increased OB–mPFC gamma connectivity [25]. Apart from symmetric connectivity metrics, directionality analyses, along with experimental approaches, have shown behaviorally-relevant information transfer between these structures [17, 21, 28, 29]. However, the roles of gamma oscillations in functional connectivity within this circuit with respect to SWM are largely unexplored.

Here, we investigated the role of the OB–mPFC circuit gamma-band activities in SWM by simultaneously recording local field potentials (LFPs) from both regions in while rats freely explored a Y-maze apparatus. We hypothesized that successful SWM performance would be associated with modulations of gamma oscillations within this circuit. We found that low gamma (30-60 Hz) power in both OB and mPFC was enhanced when the SWM was successful, and it was correlated with the animals’ behavioral performance. Also, the OB–mPFC functional connectivity was increased in both low and high gamma (60-100 Hz) sub-bands during correct trials. Granger causality (GC) demonstrated asymmetric information flow from mPFC to OB in both gamma sub-bands. Finally, we observed that phase-amplitude coupling (PAC) between mPFC delta phase and OB low gamma amplitude was enhanced during correct trials, consistent with the top-down cognitive control. Together, these findings reveal finely-tuned roles for the OB–mPFC circuit gamma oscillations in SWM.

## 2. Methods

### 2.1. Animals

Ten male Wistar rats (age: 8-10 weeks, weight: 200-250 gr) were obtained from the Laboratory Animals Research Center at Zahedan University of Medical Sciences (Zahedan, Iran). All procedures were performed consistent with the National Research Council’s Guide for the Care and Use of Laboratory Animals and the ARRIVE 2.0 guidelines. Study protocols and procedures were approved by the Ethics Committee of Zahedan University of Medical Sciences (IR.ZAUMS.AEC.1404.001). The animals were housed under standard condition (temperature: 21 ± 2 °C, 12-hour light/dark cycle, free access to water and food).

### 2.2. Surgery and electrophysiological recordings

Surgical procedures were similar to our previous studies [17, 21]. Briefly, animals first received general anesthesia by intraperitoneal injection of ketamine (100 mg/kg) and xylazine (10 mg/kg) and were placed in a stereotaxic frame (Narishige, Japan). Then, two stainless-steel recording electrodes (127 µm diameter; A.M. Systems Inc., USA) were implanted into the left OB (coordinates: AP −8.5 mm, ML −1 mm, DV −1.5 mm) and prelimbic PFC (coordinates: AP +3.2 mm, ML −0.6 mm, DV −3.6 mm) for electrophysiological recording. A stainless-steel screw was inserted into right parietal bone as the reference. All electrodes were connected to a socket, stabilized with dental acrylic cement. After post-operative care (buprenorphine injection (0.1 mg/kg) for analgesia and sterile saline injection (1.0 ml) for hydration), the animal was returned to home cage for recovery. Histological verification of electrode locations was done as previously explained [17, 21]. See our previous studies [17, 21] for further details on the experimental procedures.

On day 7 of recovery, real-time simultaneous LFPs were obtained from OB and mPFC as the animals performed the behavioral test, i.e., Y-Maze task (YMT). Throughout the recordings, head-mounted socket was connected to a miniature, high-input impedance buffer head-stage (BIODAC-A, TRITA Health Tec. Co., Tehran, Iran), that was linked via cables to a primary AC-coupled amplifier (gain: ×1000) and the data acquisition system (BIODAC-Bi401l9B, TRITA Health Tec. Co., Tehran, Iran). LFPs from OB and mPFC were recorded simultaneously with a sampling rate of 2 kHz (with 250 Hz low-pass filter) and were stored for offline analyses. Signal processing, data analyses, and visualizations were performed in Matlab 2019b (The MathWorks Inc., USA) and Python v3.8.20.

### 2.3. Behavioral procedure

After the habituation phase, the recording sessions were performed in two separate sections. First, for baseline recording, the animals were placed in their home cage and simultaneous LFPs were recorded for 10 min. Next, the recording was made as the animals were freely exploring the Y-maze apparatus, which consisted of three identical arms (50 cm in length, 10 cm in width, and 25 cm-high walls) arranged at 120°, for 10 min. Sequential transitions with consecutive visiting of all three arms were considered as correct trials (Fig. 1B). A video camera was used to record the animals’ behavior.

**Figure 1.**
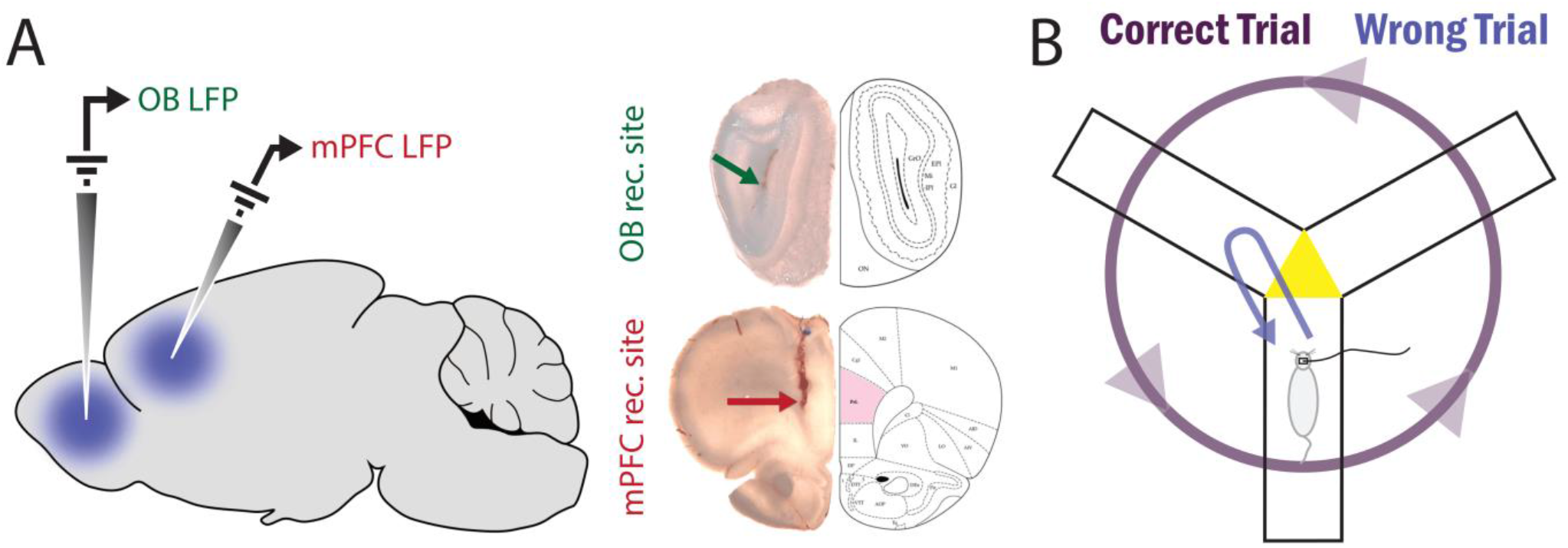
Experiment. (A) Schematic (left panel) and histological verification of OB (right upper panel) and mPFC (right lower panel) electrode sites. (B) Schematic of the WM task, i.e., YMT. Purple and blue trajectories illustrate correct and wrong trials, respectively. Adopted and modified from [17].

### 2.4. Neural data analyses

We considered 5-sec moving periods of home cage exploration and every transition in the middle triangle of Y-maze (yellow triangle in Fig. 1B) as baseline (n = 120) and task (ncorrect = 82, nwrong = 73) trials, respectively. Power spectral density (PSD) of the LFPs was estimated by convolving complex Morlet wavelets with the signals (30-160 Hz, step of 1 Hz). Normalized power values were computed by z-scoring raw power values of each trial at each frequency to the baseline for each animal separately. Average power values across time entered subsequent analyses.

We also computed magnitude-squared coherence between OB and PFC signals in different conditions (using Matlab ***mscohere*** function). Information transfer between OB and mPFC was quantified using GC [43, 44], as suggested earlier [21, 45]. For that, after down-sampled LFP signals from 500 Hz, GC for both OB-to-mPFC and mPFC-to-OB directions was computed with the model order 15, as previously suggested [21, 45]. GC values were then baseline-corrected (proportional to the baseline GC). To compute the OB–mPFC phase lag, the phase values of each region was computed by finding the angle of the analytical signal of the wavelet transform. Subsequently, phase differences averaged across time and statistically compared to 0 and π.

To compute PAC, LFP phase values were extracted using Hilbert transform and were divided into 18 phase bins (20° each). In each phase bin, the average signal amplitude was computed using Fourier transform. Subsequently, the magnitude of the mean vector that derived from the phase-binned amplitude values was defined as PAC. This vector length counted as the modulation index for subsequent statistical comparisons. To remove between-subject differences, average baseline PAC values were subtracted from each trial.

### 2.5. Statistical analyses

Statistical analyses were performed in Python v3.8.20, using scikit-learn v1.3.0 and SciPy v1.10.1 libraries, or Matlab 2019b. Circular data statistics were performed with CircStat toolbox [46] in Matlab. Details of the statistical tests are described throughout the text. Spearman correlation values between the animals’ behavioral performance and oscillatory features were computed by Matlab ***corr*** function. Permutations were repeated 10001 times.

## 3. Results

### 3.1. Simultaneous recording of OB and mPFC neural activity during SWM performance

To investigate the neural dynamics underlying SWM, we simultaneously recorded LFPs from the OB and the prelimbic region of mPFC in freely moving rats performing a YMT (Fig. 1). Trials were classified as correct when the animals successfully visited all three arms consecutively (purple trajectory in Fig. 1B), or as wrong when otherwise (Fig. 1B).

### 3.2. Regional gamma oscillations in OB and mPFC are enhanced by successful SWM performance

To characterize the oscillatory dynamics associated with SWM, we examined gamma-band activity in both OB and mPFC during correct and wrong trials. PSD analysis revealed that low gamma power was elevated during correct trials, compared to wrong trials in both OB (permutations test; Fig. 2A and Fig. 2B, left panel) and mPFC (permutations test; Fig. 2D and Fig. 2E, left panel). However, we did not find the modulation of OB (permutations test; Fig. 2A and Fig. 2B, right panel) or mPFC (permutations test; Fig. 2D and Fig. 2E, right panel) high gamma power by SWM performance.

**Figure 2.**
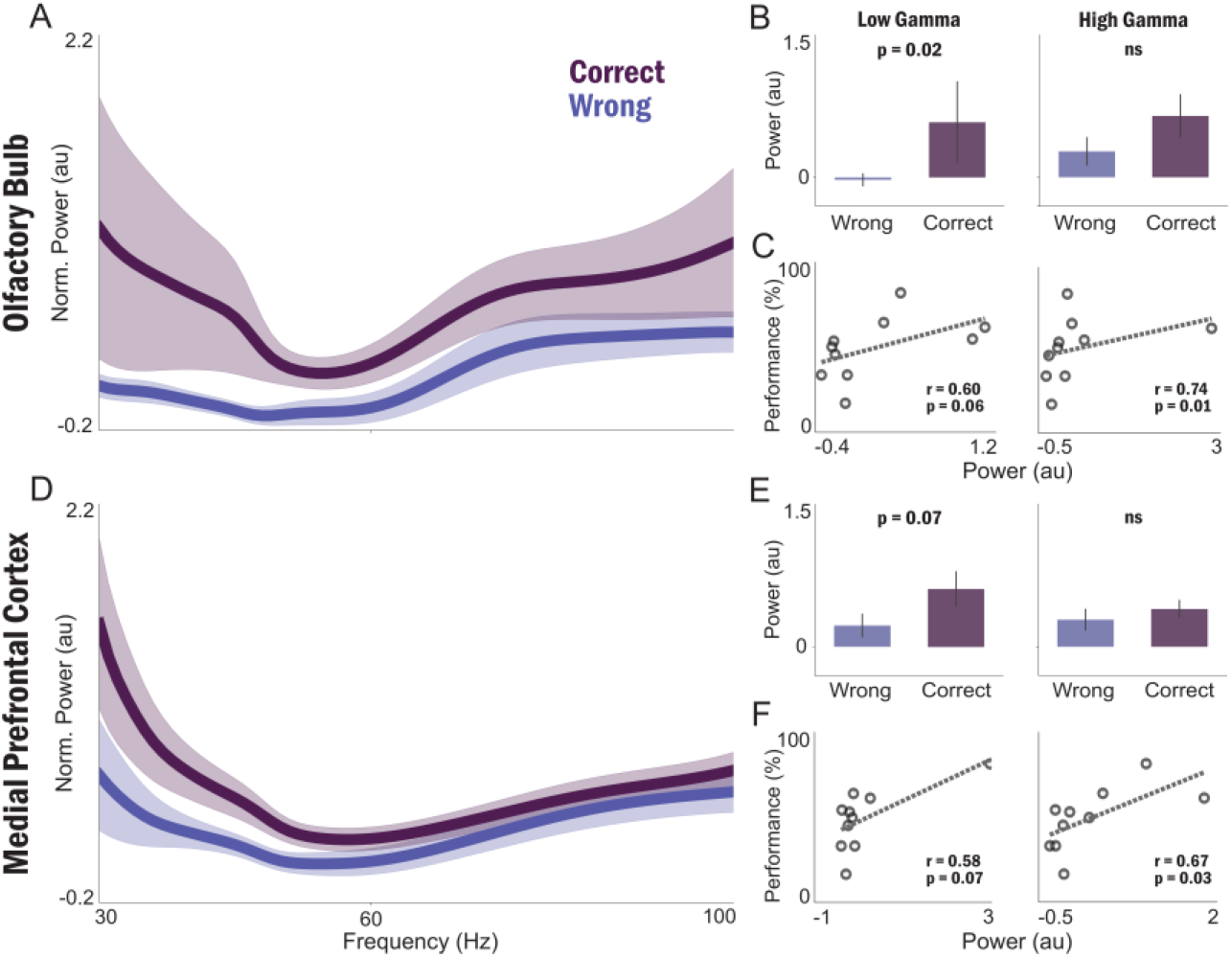
OB and mPFC regional gamma oscillations during WM. (A,D) Normalized PSD of OB (A) and mPFC (D) during correct and wrong trials. Solid line and shaded area denote mean and SEM of power, respectively. (B,E) Statistical comparison of average low (left panels) and high (right panels) gamma power of OB (B) and mPFC (E) between correct and wrong trials. Bars and error bars represent mean and SEM, respectively. (C,F) Correlation of low (left panels) and high (right panels) gamma power with behavioral performance for OB (C) and mPFC (F).

To further assess the relationship between gamma activity and SWM, we computed the correlations between gamma power and the animals’ behavioral performance. Low gamma power in both OB (spearman correlation; Fig. 2C, left panel) and mPFC (spearman correlation; Fig. 2F, left panel) exhibited positive correlations with task performance, indicating that higher low gamma power was associated with better SWM accuracy. High gamma power also showed positive correlations with performance in both regions (spearman correlation; Fig. 2C,F, right panels). Collectively, these results demonstrate that OB and mPFC gamma oscillations, particularly in the low gamma range, are dynamically linked to SWM performance.

### 3.3. OB–mPFC functional connectivity is enhanced during successful SWM performance

Having established that local gamma oscillations in OB and mPFC are modulated by SWM accuracy, we next examined whether inter-regional communication between these structures is also task-dependent. We quantified functional connectivity using spectral coherence, a measure of phase consistency between simultaneously recorded signals across frequency bands. Coherence analysis revealed robust gamma-band coupling between OB and mPFC, with distinct profiles between conditions (Fig. 3A). Statistically, both low (permutations test; Fig. 3B, left panel) and high (permutations test; Fig. 3B, right panel) gamma coherence were significantly higher during correct trials compared to wrong trials. Correlation analysis further revealed that OB–mPFC coherence in both frequency bands was positively associated with behavioral performance. Both low (spearman correlation; Fig. 3C, left panel) and high (spearman correlation; Fig. 3C, right panel) gamma coherence were positively correlated with the animals’ task performance. These results suggest that stronger inter-regional gamma synchronization is a neural signature of more accurate SWM processing.

**Figure 3.**
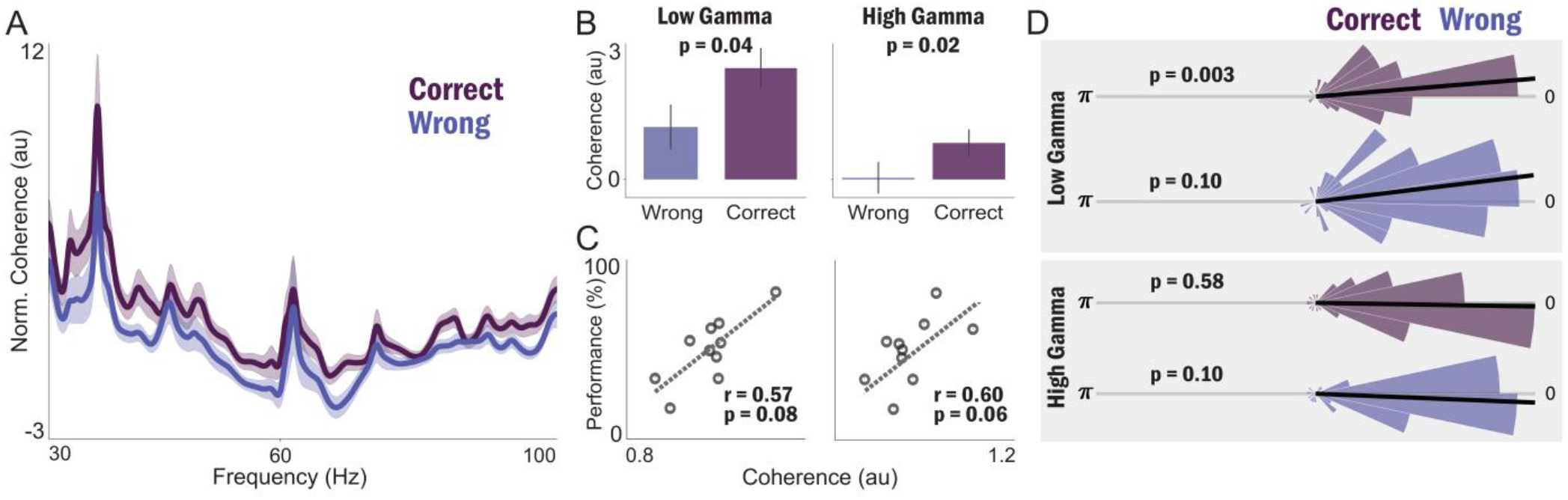
OB–mPFC functional connectivity across gamma oscillations during WM. (A) Normalized OB–mPFC gamma oscillations spectral coherence. Solid line and shaded area denote mean and SEM of coherence, respectively. (B) Statistical comparison of average OB–mPFC coherence in low (left panel) and high (right panel) gamma bands between correct and wrong trials. Bars and error bars represent mean and SEM, respectively. (C) Correlation of OB–mPFC coherence in low (left panel) and high (right panel) gamma bands with behavioral performance. (D) Polar distribution of OB–mPFC phase-lag in low (upper two panels) and high (lower two panels) gamma bands for correct and wrong trials. Black line indicates median phase lag across trials. Statistical comparison with binomial test from zero rad.

Brain areas typically communicate through synaptic connections, i.e., synaptic information transfer; yet the detection of enhanced functional connectivity could also be due to volume-conduction [45]. In the case of synaptically-conducted connectivity, the phase lag between the two time-series differs from zero and π. Here, we found that the phase lag between OB and mPFC low gamma differed from zero and π in correct trials, but not in wrong trials (binomial test; Fig. 3D, upper panel). In high gamma oscillations, phase lag distributions did not deviate significantly from zero and π in either trial type (binomial test; Fig. 3D, lower panel). Therefore, overall, we show that that the OB–mPFC gamma band functional connectivity, especially in the low gamma range, is enhanced during SWM, most probably to support synaptic information transfer.

### 3.4. Directional information flow from mPFC to OB supports successful SWM performance

To characterize the directionality of information transfer between OB and mPFC, we computed GC, which quantifies the extent to which past activity in one signal predicts future activity in another [43, 44], across the gamma frequency range. In the mPFC → OB direction, GC was significantly elevated during correct, compared to wrong, trials in both low (permutations test; Fig. 4A and Fig. 4B, left panel) and high (permutations test; Fig. 4A and Fig. 4B, right panel) gamma bands. These findings indicate that top-down information flow from mPFC to OB supports the animals’ SWM performance.

**Figure 4.**
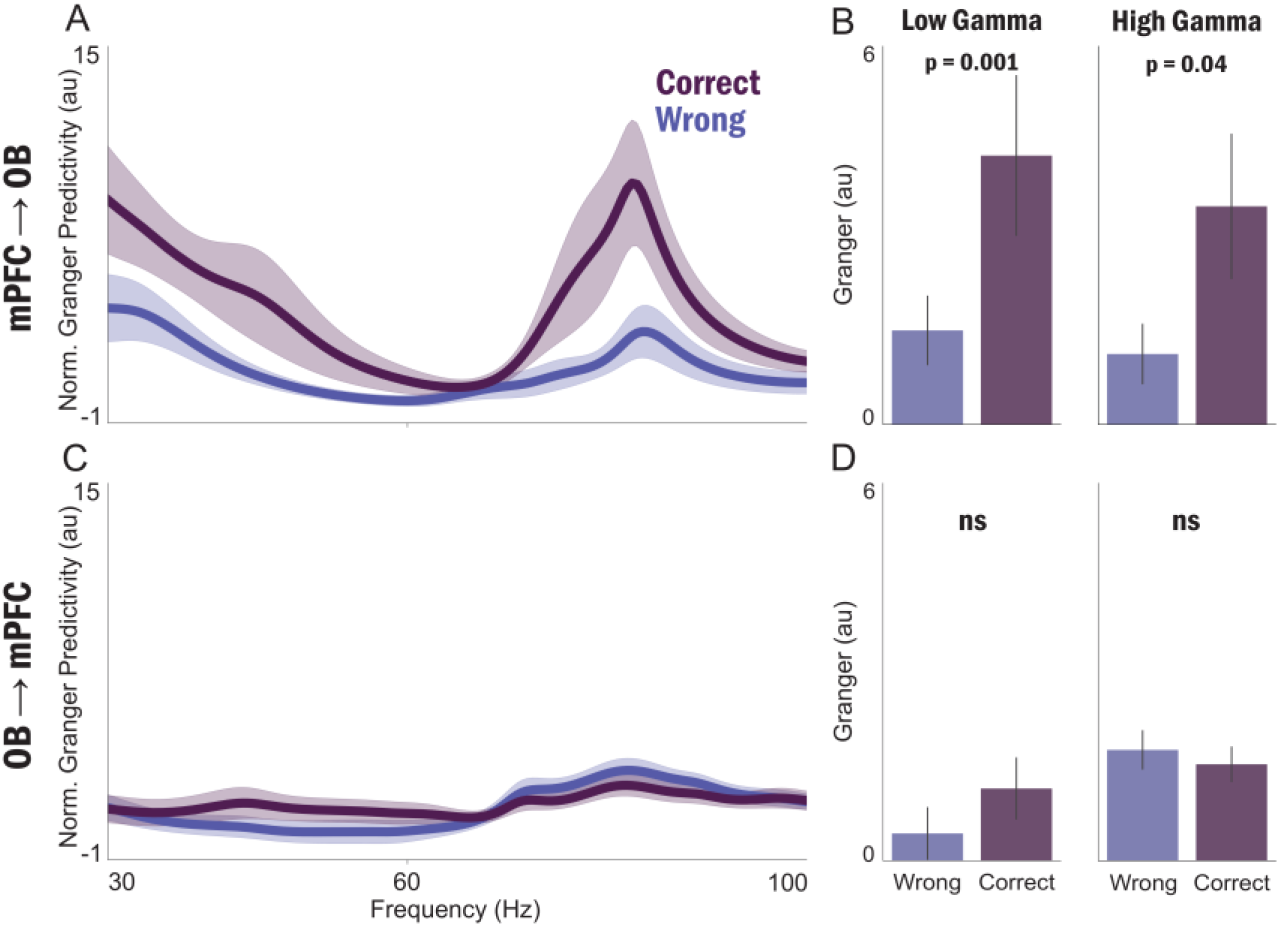
Gamma band information transfer between OB and mPFC during WM. (A,C) Normalized GC in the mPFC-to-OB (A) and OB-to-mPFC (C) directions in correct and wrong trials. Solid line and shaded area denote mean and SEM, respectively. (B,D) Statistical comparison of average GC in low (left panels) and high (right panels) gamma bands between correct and wrong trials in the mPFC-to-OB (B) and OB-to-mPFC (D) directions. Bars and error bars represent mean and SEM, respectively.

In contrast, for the opposite direction, i.e., OB → mPFC, the GC was generally weaker, particularly in correct trials, and we found no significant GC difference between correct and wrong trials in either gamma band (permutations test; Fig. 4C,D). This absence of performance-dependent modulation in the bottom-up direction suggests that feedforward signaling from OB to mPFC remains relatively stable regardless of trial outcome.

### 3.5. mPFC–OB delta-gamma phase-amplitude coupling supports successful SWM performance

Beyond spectral coherence and GC, we investigated whether cross-frequency coupling between regions contributes to SWM performance. Specifically, we examined inter-regional PAC, which evaluates the modulation of one region’s amplitude by another region’s slow oscillations phase. We computed modulation index (MI) for both mPFC–OB and OB–mPFC PAC, testing delta and theta phase coupling with low and high gamma amplitudes, separately for correct and wrong trials.

For mPFC–OB PAC, we observed enhanced coupling between mPFC delta phase and OB low, but not high, gamma amplitude during correct, compared to wrong trials (permutations test; Fig. 5A,B,D and Fig. 5C,E, left panels). Also, the mPFC–OB delta-low gamma PAC was correlated with the animals’ behavioral performance (spearman correlation; r = 0.60, p = 0.055; not shown). However, we did not find a significant difference in the mPFC–OB theta-gamma PAC between correct and wrong trials (permutations test; Fig. 5A,B,D and Fig. 5C,E, right panels); nor did we find any differences in the OB–mPFC PAC (permutations test; Fig. 5F-J). These findings demonstrate that inter-regional cross-frequency coupling is highly selective: only mPFC delta phase coupling with OB low gamma amplitude is enhanced during successful SWM performance. This specific pattern of mPFC–OB delta-low gamma PAC along the low gamma GC (see Fig. 4), suggests that prefrontal feedbacks coordinate OB faster oscillations to enhance the animals’ SWM processing.

**Figure 5.**
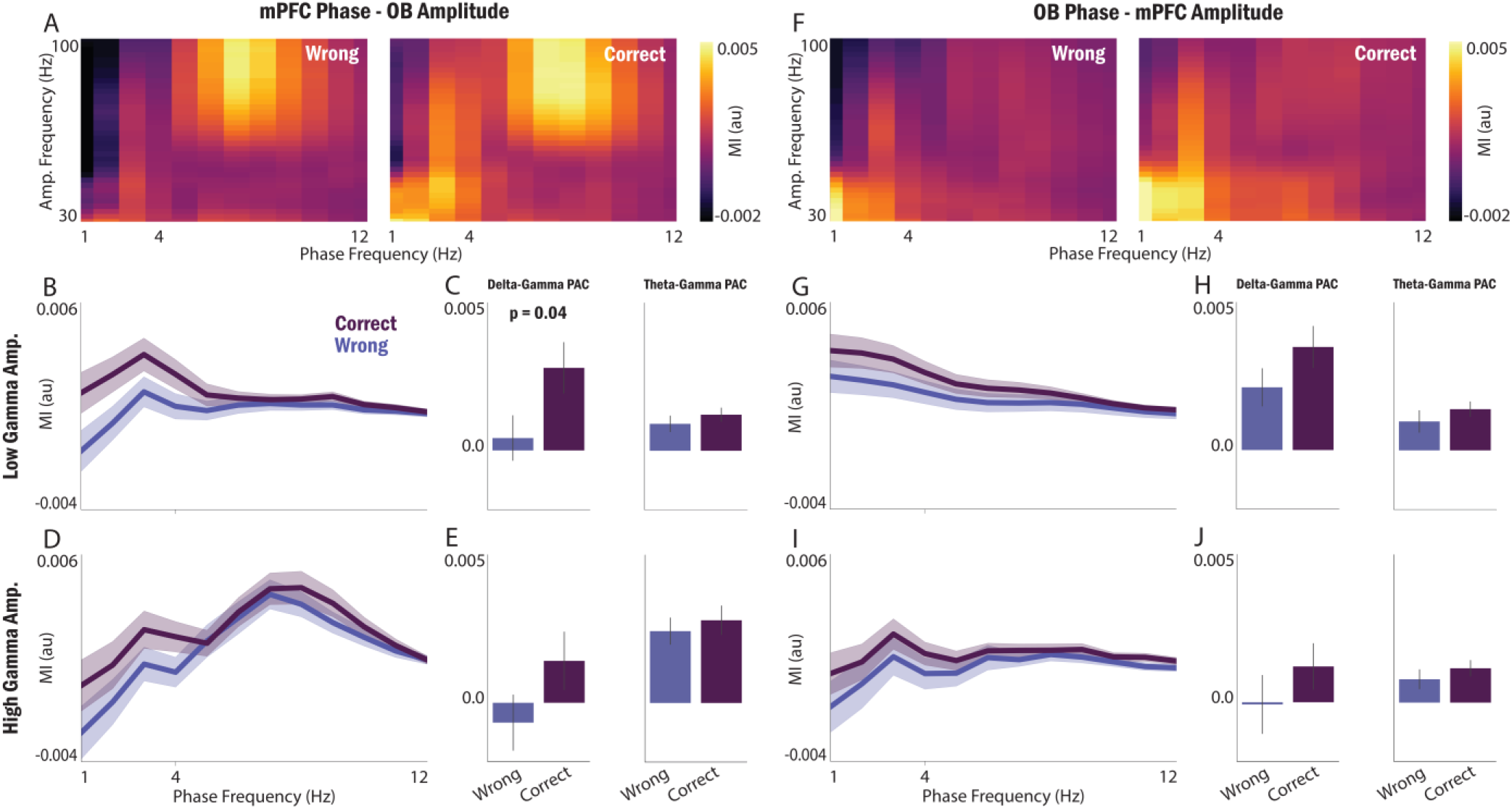
Inter-regional delta- and theta-gamma PAC between OB and mPFC during WM. (A,F) Average of normalized MI for mPFC–OB (A) and OB–mPFC (F) PAC in correct (right panels) and wrong (left panels) trials. (B,D,G,I) Normalized MI of mPFC–OB (B,D) and OB–mPFC (G,I) PAC for low (B,G) and high (D,I) gamma amplitudes during correct and wrong trials. Solid line and shaded area denote mean and SEM, respectively. (C,E,H,J) Statistical comparison of average delta (left panels) and theta (right panels) phase PAC for low (C,H) and high (E,J) gamma amplitudes during correct and wrong trials. Bars and error bars represent mean and SEM, respectively.

## 4. Discussion

The present study characterizes the oscillatory dynamics in the OB–mPFC circuit during SWM, using simultaneous recordings in freely behaving rats performing a YMT. Specifically, we find that successful SWM: (1) elevates low gamma power in both OB and mPFC in correlation with behavioral performance; (2) enhances inter-regional gamma band functional connectivity, most probably through synaptic mechanisms; (3) improves top-down feedbacks as evidenced by enhancement of mPFC-to-OB information flow in both gamma sub-bands and mPFC–OB delta-low gamma PAC. Together, these findings reveal that successful SWM is supported by coordinated gamma-band oscillations, enhanced functional connectivity, and top-down directional control mechanisms within the OB–mPFC circuit.

Prefrontal and sensory cortical gamma activity has long been associated with the maintenance of representations and encoding of task-relevant information [1, 33, 38-40]. Furthermore, gamma oscillations are shown to support cognitive functions, such as WM, attention, and decision-making, through transient coordination of distributed neural assemblies [1, 16, 35, 37-40, 47-49]. Our results demonstrate that this gamma-mediated coding mechanism is not restricted to mPFC but is also present in the OB, a region traditionally viewed as a primary sensory relay [18-20]. Furthermore, the selective enhancement of low, but not high, gamma power during SWM suggests that these two gamma sub-bands may reflect distinct computational processes. The correlation between low gamma power and performance further supports the hypothesis that gamma oscillations carry behaviorally relevant information and may serve as a neural substrate for the precision and stability of SWM representations.

Beyond local oscillatory changes, we observed enhanced coherence between OB and mPFC in both low and high gamma bands during successful SWM. Coherence reflects the temporal coordination of neural activity across distant brain regions and is thought to facilitate information integration and communication [14, 21, 24, 26, 29, 30, 32, 33, 50]. Our findings are consistent with the “communication through coherence” framework, which posits that synchronized oscillations, especially gamma-band connectivity, may act as communication channels between neuronal populations, enabling effective transmission of information [33]. The performance-dependent enhancement of gamma coherence during SWM suggests that successful SWM performance requires not only local processing within each region, but also coordinated activity across the OB–mPFC circuit. Moreover, because of the fast temporal dynamics of gamma rhythms, these oscillations might enable rapid neural coordination and information transfer between distant populations. From another perspective, the OB is not traditionally viewed as a canonical SWM structure; however, accumulating evidence indicates that it participates in large-scale brain networks and can coordinate with prefrontal and hippocampal regions during cognitive and emotional behaviors [12, 18, 21-29, 42, 51, 52]. The present findings support the view that OB oscillatory activity is not merely sensory, but can be recruited into broader networks supporting a range of behaviors. These results further extend previous reports of OB–mPFC connectivity during anxiety and fear-related behaviors [21, 22, 28, 29] to the cognitive domain, supporting the hypothesis that the OB and mPFC form a flexible, and probably domain-general (see [21] for a discussion on this issue), circuit for both emotional and cognitive functions.

The prefrontal cortex is known to exert top-down influence over sensory regions to guide goal-directed behavior, suppress irrelevant information, and maintain task-relevant representations [1-3, 9, 10, 53, 54]. Beyond functional connectivity, our results reveal directional information flow from mPFC to OB in the gamma frequency range (see Fig. 4). Consistent with previous evidence [17], these findings suggest that the top-down cognitive control theory extends to the OB as well, particularly in the present case of SWM. Such directionality is consistent with the general role of mPFC in executive functions. The directionality of information flow in our study, i.e., mPFC → OB, suggests that the prefrontal cortex may use gamma-band signaling to coordinate or “instruct” OB activity, possibly to suppress task-irrelevant sensory input or to enhance task-relevant representations. In the context of SWM, the mPFC may integrate ongoing spatial information, and then bias, in other words modulate, the OB oscillatory activities to support successful target selection and sequential navigation. The fact that gamma-band GC was enhanced in the mPFC → OB direction suggests that prefrontal feedback may be a key mechanism by which cognitive demands are translated into coordinated network dynamics.

Furthermore, we show that the phase of mPFC delta oscillations modulates the OB gamma activity, a phenomenon which is more prominent during successful SWM performance (see Fig. 5). PAC is thought to reflect a hierarchical organization of neural oscillations, in which slow rhythms coordinate the timing of faster rhythms and thereby facilitate information integration across spatial and temporal scales as well as coordinate local processing with inter-regional exchange [55, 56]. In the context of SWM, delta oscillations may provide a temporal framework that organizes gamma-band activity into discrete processing windows, enabling the sequential encoding, maintenance, and retrieval of information. Our finding that PAC is stronger during successful SWM suggests that the temporal coordination of slow and fast rhythms is a behaviorally relevant mechanism that supports successful SWM performance. Furthermore, this PAC was observed between mPFC delta phase and OB gamma amplitude, suggesting a feedback mechanism and further signifying the roles mPFC modulations over OB oscillations during SWM. Together with our coherence and GC findings, the PAC results support a model in which mPFC oscillations orchestrate OB gamma-band activity to support WM, most probably by filtering-out task-irrelevant information from the sensory input. Along others [57], we propose that mPFC slow oscillations serve as a “master clock” that coordinates gamma-band activity across the circuit, while gamma oscillations carry the content of WM.

Our findings contribute to a growing body of evidence that the OB plays a broader role in cognition and emotion than previously appreciated [23-28]. While the OB has traditionally been studied in the context of odor processing [18, 20], recent work has implicated OB oscillations in anxiety, fear, and recognition memory [21, 22, 24-26, 28, 29]. Anatomical studies reveal that the OB is connected not only to primary olfactory cortex, but also to mPFC, hippocampus, and so on [42]. Our results extend this literature by demonstrating that the OB–mPFC circuit is functionally engaged during SWM. Furthermore, we support the idea that OB is not merely a passive relay, but an active participant in the neural computations underlying a range of behaviors. This interpretation is consistent with recent proposals that sensory regions contribute to cognitive control in ways such as providing contextual information, maintaining sensory representations, and predictive coding [3, 53, 54, 58, 59].

The present study has specific limitations that should be acknowledged. Our analyses were based on LFP recordings, which reflect the summed activity of neuronal populations and do not provide information about the spiking activity of individual neurons. Future studies should combine LFP recordings with single-unit recordings to determine how oscillatory dynamics relate to the firing patterns of specific cell types and how neuronal ensembles encode SWM representations. Second, while we observed robust associations between gamma power, coherence, and performance, causal perturbation studies using optogenetics, chemogenetics, or closed-loop stimulations are needed for establishing to what extent the OB–mPFC circuit gamma oscillations affect SWM performance. Finally, future studies should incorporate respiration monitoring to determine whether respiration-related rhythms contribute to OB–mPFC gamma activities during SWM.

Taken together, our results suggest that SWM is supported by multiple complementary mechanisms operating within the OB–mPFC circuit. First, local gamma oscillations in both regions encode task-relevant information. Second, inter-regional coherence in the gamma range reflects synchronized activity that may facilitate information transfer and integration. Third, directional information flow from mPFC to OB introduces a neural channel for top-down cognitive control. Fourth, cross-frequency coupling between mPFC delta phase and OB gamma amplitude provides a temporal framework for organizing OB neural activity. These mechanisms are not mutually exclusive but likely operate in concert to support the encoding, maintenance, and retrieval of spatial information during SWM. Top-down signaling from mPFC to OB may serve to align OB activity with task demands, suppress irrelevant sensory input, or enhance task-relevant representations. This multi-mechanism model is consistent with recent theoretical frameworks emphasizing the importance of distributed neural coding, hierarchical oscillatory organization, and flexible inter-regional communication in cognitive control [2, 10, 11, 53, 54]. These findings support the idea that the OB is not merely a sensory relay but an active participant in cognitive processing, and that the OB–mPFC circuit serves as a flexible communication channel for both cognitive and emotional functions.

## Author Contributions

MM conceptualized the study. MM and ASM performed the surgeries, ran the experiments, collected the data, designed the analysis plan, analyzed the data, and drafted the manuscript. All authors reviewed the manuscript. MRR and MAM supervised the study.

## Acknowledgements

The authors would like to thank Farhad Tabasi and Morteza Salimi for their constructive comments on the experimental design and procedures. We also appreciate technical assistance from Soheil Karimi Darmian.

## Data and Code Availability

Data, Matlab codes, and Python notebooks will be made available upon reasonable request to the corresponding author.

## Funding

This work was funded by Zahedan University of Medical Sciences (grant number: 11707).

## Competing Interests

None declared.

## Notes

### Competing Interest Statement

The authors have declared no competing interest.

